# An iteratively curated CRISPR library reveals target-specific biological resistance landscapes across targeted protein degraders

**DOI:** 10.64898/2026.08.14.744749

**Authors:** Lin Liu, Olivia Voulgaris, Changqing Wang, Dáire Gannon, Matthew E Ritchie, Rebecca Feltham, Stephin J. Vervoort

## Abstract

Targeted protein degradation (TPD) has emerged as an increasingly powerful approach for therapeutic development and biological discovery. TPD compounds including proteolysis-targeting chimeras (PROTACs), molecular glues, and tag-targeting protein degraders (tTPD) enable rapid, selective and reversible degradation of proteins through recruitment of the ubiquitin-proteasome system (UPS). However, genome-wide CRISPR screens performed with targeted protein degraders are frequently dominated by resistance mechanisms that disrupt degrader activity, including loss of recruited E3 ligase components and broader UPS regulators. The strong selective advantage conferred by these perturbations can obscure less penetrant, biological genetic interactions that operate downstream of target degradation. To overcome this limitation, through iterative genome-wide screening and manual curation, we developed a TPD-compatible CRISPR knockout library that retains near-genome-scale coverage while excluding a focused set of genes recurrently associated with degrader failure. Across multiple degrader screens, this library reduced the dominance of UPS-associated resistance mechanisms and improved the detection and prioritization of genetic interactions linked to target biology. Using the RBM39 molecular glue degrader indisulam as a model, we identified ZMAT2 loss as a resistance mechanism that preserves RBM39 degradation but attenuates the transcriptional and splicing consequences of target depletion. Together, our work establishes a TPD-compatible CRISPR screening framework that improves the biological resolution of degrader resistance screens and facilitates the discovery of genetic dependencies operating downstream of targeted protein degradation.

## Introduction

Targeted protein degradation (TPD) uses small molecules to induce the selective degradation of target proteins^1^. While genetic tools such as CRISPR-Cas9 knockout are powerful, they result in permanent protein loss, which may lead to compensatory effects. Traditional small-molecule inhibitors allow rapid perturbation, yet they are limited by off-target effects. Furthermore, small-molecule inhibitors typically require sustained occupancy of an active or allosteric binding site, rendering non-enzymatic and scaffolding proteins that lack ligandable active sites difficult to target. Targeted protein degradation enables rapid, acute, precise and reversible protein depletion, presenting exciting opportunities to explore the previously undruggable proteome^2^.

Most TPD strategies trigger intracellular protein degradation by hijacking the ubiquitin-proteasome system (UPS)^3^, which catalyses protein ubiquitination through an enzymatic cascade involving E1 activating enzymes, E2 conjugating enzymes, and E3 ubiquitin ligases, prompting subsequent proteasomal degradation^4^. A major type of E3 ligases currently exploited by TPD belong to the cullin-RING ligase (CRL) family. CRL complexes are multisubunit E3 ligases composed of a cullin scaffold, a RING finger protein that recruits the E2 enzyme, a variable substrate receptor (SR; e.g., Cereblon (CRBN), VHL, or DCAF15), and an adaptor protein that links the SR to the cullin scaffold (e.g., DDB1, SKP1, or Elongin B/C)^5^. The CRL complexes are activated by the conjugation of the ubiquitin-like protein NEDD8, while deNEDDylation by the COP9 signalosome (CSN) inactivates the complexes^6^.

The two major classes of TPD small-molecule degraders that rely on the ubiquitin-proteasome system are proteolysis-targeting chimeras (PROTACs) and molecular glues^7^. PROTACs are heterobifunctional molecules composed of a protein of interest (POI)-binding ligand and an E3 ligase-recruiting ligand connected by a chemical linker^2^. In contrast, molecular glue degraders are monovalent small molecules that stabilize or facilitate interactions between an E3 ligase and a target protein by reshaping their binding interface, thereby facilitating ubiquitylation and subsequent degradation through the UPS^8^.

Given their unique advantages over conventional protein inhibition, TPD has emerged as a promising therapeutic strategy. The first clinical proof of concept is provided by the immunomodulatory drugs (IMiDs) thalidomide, lenalidomide, and pomalidomide, which were later recognized as molecular glue degraders^9^. The anticancer aryl-sulfonamide indisulam (also known as E7070) functions as a molecular glue degrader by promoting the interaction between pre-mRNA splicing factor RNA-binding motif protein 39 (RBM39) and the CRL4-DCAF15 E3 ubiquitin ligase, leading to proteasomal degradation of RBM39 and splicing defects^10, 11^. Several PROTACs targeting the Bromodomain and Extra-Terminal (BET) family proteins, such as dBET6^12^, MZ1^13^ and ARV771^14^ have shown potent antitumor efficacy. More recently, the heterobifunctional vepdegestrant (also known as ARV471)^15^, an estrogen receptor (ER) degrader, became the first PROTAC approved by the US Food and Drug Administration, for adults with previously treated, ESR1-mutant, ER-positive/HER2-negative advanced or metastatic breast cancer^16^.

Alongside chemical degraders, tag-targeting protein degraders (tTPD) or degron tags, which include PROTAC– or molecular glue-based compounds, provide a generalizable approach to rapidly and selectively degrade allele-specific protein and are now extensively used for biological studies ^7, 17, 18^. Common degron tags include the FKBP12^36V^ Tag ^19^, BromoTag ^20^, HaloTag ^21^ and the monovalent molecular glue-based auxin-inducible degron (AID Tag) ^22^. Upon addition of a cell-permeable, tag-specific degrader, the degron tagged protein is recruited to a cullin-RING E3 ubiquitin ligase complex, resulting in its ubiquitylation and proteasomal degradation.

High-throughput functional genomic approaches, including RNA interference and genome-wide CRISPR-Cas9 knockout screens, have become powerful tools for identifying mechanisms of essentiality, drug resistance mechanisms, and therapeutic vulnerabilities ^23, 24, 25, 26^. Recently, these screening strategies have been combined with targeted protein degradation to investigate the biological consequences of acute protein loss ^18^. However, genome-wide CRISPR screens using PROTACs or molecular glue degraders have consistently identified components of the UPS machinery as the dominant resistance hits, which include the recruited E3 ligase complex, E2 ubiquitin-conjugating enzymes, neddylation factors, and the COP9 signalosome ^12, 27^. Disruption of the UPS degradation pathway abolishes target protein degradation irrespective of its biological function, conferring near-complete resistance and a strong selective advantage. Consequently, clones carrying defects in the UPS degradation machinery rapidly outcompete other resistant populations, masking weaker, but true biological genetic interactors of target protein. These observations suggested that conventional genome-wide CRISPR screening is poorly suited for uncovering target-specific biological determinants using TPD compounds, highlighting the need for TPD-compatible screening strategies that uncouple degradation resistance from target-dependent biological mechanisms.

Here we set out to create a CRISPR knockout-library compatible with TPD and tag-TPD systems to enable biological discovery. To this end, we systematically examined the resistance landscape across degraders targeting distinct proteins and recruiting multiple E3 ligases. We reasoned that selectively removing UPS-mediated resistance genes would improve the recovery of mechanisms operating downstream of target depletion. Through iterative genome-wide screening and manual curation, we developed a TPD-compatible CRISPR knockout library that retains near-genome-scale coverage while excluding a focused set of genes associated with degrader failure. Using the molecular glue indisulam as a model, we further identify ZMAT2 loss as a resistance mechanism that preserves RBM39 degradation but buffers the transcriptional and splicing consequences of RBM39 loss.

## Results

### UPS-dominated genome-wide CRISPR screens with TPD compounds lead to the development of a TPD-compatible CRISPR library

The activity of heterobifunctional degraders, molecular glues, and tag-targeting protein degraders (tTPD) such as dTAG ^19^, is reliant on target engagement in conjunction with efficient recruitment of the ubiquitin-proteasome (UPS) system. Previous genome-wide CRISPR screens of targeted protein degraders (TPDs) have shown that resistance landscapes are dominated by genes that are part of the UPS system, the inactivation of which prevents efficient target degradation. Potent resistance genes include the TPD engaged E3 ligases, such as Cereblon, VHL and DCAF15, other E1 and E2 components as well as associated machinery such as the COP9 signalosome ^12, 27^.

In a pooled CRISPR screen, each clone ideally carries a defined genetic perturbation and undergoes exponential growth within a closed cell population, where clones compete with one another based on their relative fitness. Screen analysis quantifies changes in the relative abundance of sgRNAs within the population rather than absolute changes in clone number ^28^. To assess how the presence of degrader-failure clones may impact the identification of biological resistance hits, we performed *in-silico* simulations of clonal dynamics in a pooled CRISPR screen containing two classes of resistance mechanisms: near-complete resistance (degrader-failure), and less penetrant resistance (biological resistance), which acts downstream of degradation. Simulated CRISPR-screening with an unfiltered library inevitably resulted in the acquisition of degrader failure clones which rapidly expand to dominate the population. As a result, at the end point the abundance of all other clones, including those possessing biological resistance, were largely compressed and exhibited minimal enrichment in pooled readouts. However, simulated CRISPR screening with a filtered library, where degrader-failure clones were removed, showed increased simulated log-fold change (LFC) of biological resistors at the simulated endpoint (Fig. 1A).

**Figure 1.**
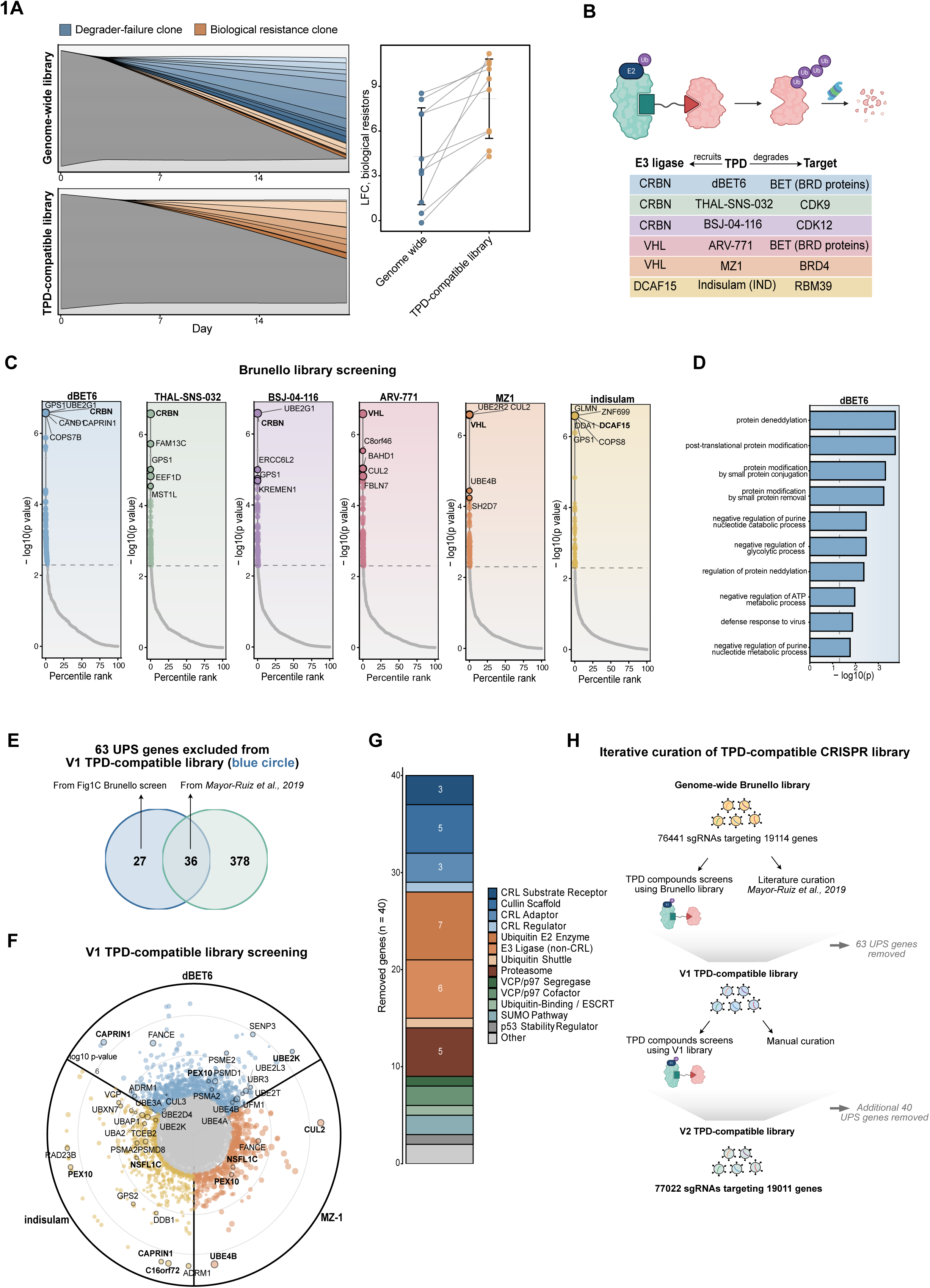
UPS-dominated genome-wide CRISPR screens prompt the development of a TPD-compatible knockout library. **(A)** *In-silico* simulation of clonal dynamics in a pooled CRISPR screen over 21 days of degrader selection. A population of 20,000 cells was partitioned into ten near-complete (degrader-failure) resistance clones (blue), ten less-penetrant biological resistance clones (orange) and a non-resistant background pool (grey). *Left*, stream plots of clone fraction over time for a genome-wide library and for a TPD-compatible library, in which the corresponding sgRNAs are absent and these clones do not expand. *Right*, simulated log2 fold change of each biological resistance clone between day 0 and day 21, computed from its fraction of the total population, in the two library scenarios. Grey lines connect the same clone across scenarios; bars show mean ± s.d. (n = 10 clones). **(B)** Schematic of targeted protein degraders (TPDs) used in this study. The recruited E3 ligases, the degraders, and the targets are indicated: CRBN–dBET6–BET (BRD proteins); CRBN–THAL-SNS-032–CDK9; CRBN–BSJ-04-116–CDK12; VHL–ARV-771–BET; VHL–MZ1–BRD4; DCAF15–indisulam–RBM39. **(C)** Ranked significance plots of the six genome-wide Brunello resistance screens using TPD compounds described in (B). Genes are ordered by percentile rank of the MAGeCK RRA positive-selection *p* value and plotted against −log10(*p*). The dashed line marks the hit threshold (*p* < 0.005); point size scales with gene-level log2 fold change. Coloured points are resistant hits (*p* < 0.005, LFC > 1). Circles with outline indicate the top 5 resistant hits. The recruited E3 ligase substrate receptors (CRBN, VHL, DCAF15) are highlighted in bold. **(D)** Gene Ontology Biological Process over-representation among significant resistance hits from the Brunello dBET6 screen (g:Profiler; Benjamini–Hochberg FDR). The ten most significant terms after removal of generic parent terms and de-duplication of near-identical labels are shown. Bars, −log10(*p*); dashed line, *p* = 0.05. **(E)** Overlap between the 63 genes excluded from the V1 library and the top 100 resistance hits per screen from *Mayor-Ruiz et al., 2019*^12^. **(F)** Circular rank plot of resistance hits from the three TPD-compatible V1 library screens (dBET6, MZ-1, indisulam; compared with DMSO). Radial position gives −log10(RRA *p*); size of the circle indicates LFC values. Circles with outline indicated the 30 V1 screen hits subsequently excluded from the V2. Hits of note are highlighted in bold. **(G)** Functional classification of genes identified for removal in V2 library. Bars are stacked by manually curated ubiquitin-proteasome system category. **(H)** Schematic of iterative curation of TPD-compatible CRISPR library.

To experimentally define the dominant resistance landscape, we performed genome-wide CRISPR screens using the Brunello library across a panel of PROTAC and molecular glue degraders spanning multiple E3 ligase complexes (CRL4-CRBN, CRL4-DCAF15, and CRL2-VHL) and targets (BRD proteins, RBM39, CDK9 and CDK12) (Fig. 1B). Across all screens, the top resistance hits corresponded to the recruited E3 ligase complexes (CRBN, VHL and DCAF15) and established regulators of CRL function (UBE2G1, GPS1 and CAND1) (Fig. 1C). Using the dBET6 screen as an example, gene ontology analysis of significant hits revealed strong enrichment for ubiquitin system-associated processes, including protein deNEDDylation, and other post-translational regulatory pathways (Fig. 1D).

In clinical settings, the identification of biological resistance may be important, as genetic and non-genetic adaptation to target engagement may arise from mechanisms beyond the UPS system. Similarly, genetic interactions related to the target biology rather than the UPS are important for molecular and cell biology studies in which tTPD systems have been widely used to study target function. Having established that the dominance of degrader-failure mechanisms can obscure biological genetic dependencies operating downstream of target degradation *in-silico*, we generated a TPD-compatible library (V1) by excluding 63 UPS genes from the Brunello library (Fig. 1E). This exclusion list included 36 UPS-related genes from literature^12^ and additional 27 genes from our TPD compound Brunello screening as shown in Fig. 1C, spanning many functions of the UPS system (Supplementary Fig. 1A).

The V1 TPD-compatible library was subsequently used in screens with representative degraders spanning three E3 ligase classes: indisulam (DCAF15), dBET6 (CRBN), and MZ1 (VHL) (Fig. 1F). Despite substantially improved detection of less penetrant resistance hits, cullin-RING ligase associated genes continued to emerge as top hits, including CRL2 (also known as CUL2) and UBE4B. The screens further recapitulated UPS-associated resistance factors identified in previous TPD screens, including CAPRIN1 and C16orf72 (HAPSTR1) ^12, 27^. CAPRIN1 functions within the CAPRIN1–CUL1–FBXO42 ubiquitin ligase complex, and C16orf72 (HAPSTR1) is required for HUWE1 nuclear localisation and substrate targeting ^29, 30^. Beyond these established factors, the V1 screens nominated previously uncharacterised resistance candidates with potential roles in degrader activity. UBE2K, a ubiquitin-conjugating (E2) enzyme containing a UBA domain that elongates K48-linked and K48/K63-branched polyubiquitin chains ^31, 32^, was recovered in the dBET6 screen. Given that CRL4–CRBN neosubstrate ubiquitination has been attributed primarily to UBE2G1 and UBE2D3^33, 34^, this nominates an additional E2 that may contribute to degrader-induced chain formation. NSFL1C (p47), a UBA-domain cofactor of the p97/VCP segregase^35^, was identified in both the MZ1 and indisulam screens, implicating the substrate-extraction step downstream of ubiquitylation. PEX10, a RING-type E3 ligase of the peroxisomal PEX2-PEX10-PEX12 import complex^36^, was recovered across all three V1 library screens despite having no established role in cullin-RING-mediated degradation.

These findings suggest that sensitivity to targeted degradation is governed by a broader ubiquitin-proteasome network extending beyond the recruited cullin-RING E3 ligase complex, spanning chain elongation, substrate extraction, and potentially non-canonical ligase activities. Building on these observations, we further refined the V1 TPD-compatible knockout library by excluding 30 additional UPS-associated genes identified across the V1 screens, together with 10 manually curated UPS genes (Fig. 1G). Through this iterative curation process, we generated the V2 TPD-compatible CRISPR library (hereafter referred to as the V2 library). Compared with the Brunello library, the final V2 library excluded a total of 103 UPS-associated genes while retaining near-genome-scale coverage, comprising 77022 sgRNAs targeting 19011 genes (Fig. 1H).

### An iteratively curated TPD-compatible CRISPR library enables the discovery of genetic interactions linked to target biology

Network visualisation of the 103 UPS-associated genes excluded from the V2 library revealed that they spanned multiple functional branches of the ubiquitin-proteasome system, including cullin-RING ligase regulation, E2/E3 ubiquitin transfer, neddylation and deNEDDylation, deubiquitination, SUMOylation, proteasome function, and broader ubiquitin pathway regulation (Fig.2 A, B).

**Figure 2.**
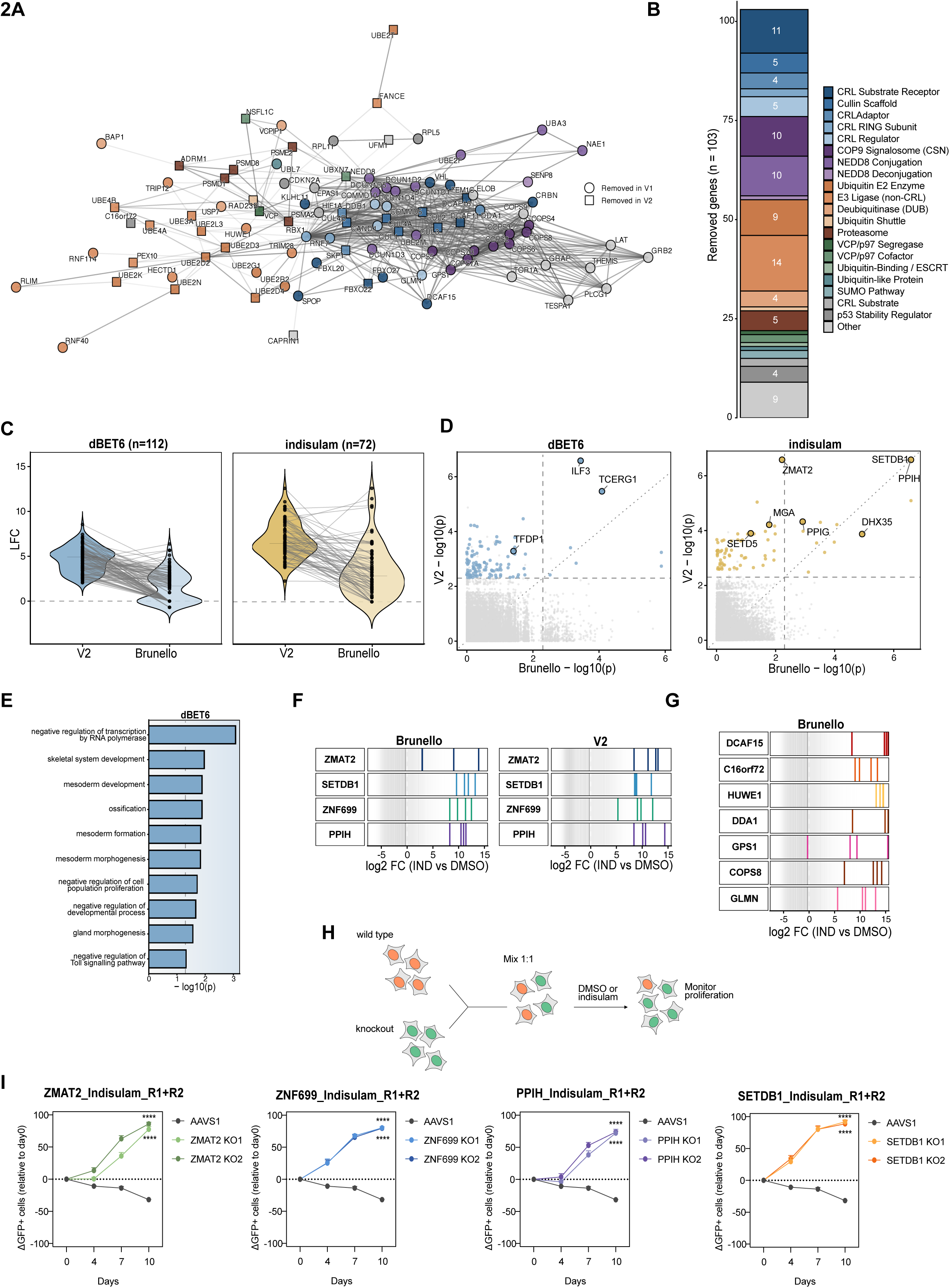
A TPD-compatible CRISPR library (V2) exposes genetic interactions linked to target biology. **(A)** Network visualisation of UPS-associated genes (n=103) excluded from the V2 library using STRING v12.0 (*Homo sapiens*). Nodes are coloured by manually curated UPS functional category; edges are STRING physical associations with thickness corresponding to confidence score. Circle nodes indicate V1 removed genes. Square nodes are nodes excluded after V1 screens. **(B)** Functional classification of the 103 excluded genes. Bars are stacked by UPS category. **(C)** Gene-level log2 fold change (dBET6 or Indisulam vs DMSO) of significant V2 resistance hits (p < 0.005, LFC > 1), compared with the log2 fold change of the same genes in the matched Brunello screen. Brunello reads were counted against the V2 reference library so that both analyses share an identical gene universe, normalisation base and sgRNA composition; *n* = number of V2 hits per degrader. Dashed line, LFC = 0. **(D)** Concordance of statistical significance between the Brunello and V2 screens for dBET6 (left) and indisulam (right). Each point is a gene; axes give −log10(RRA positive-selection *p*) in each library. Dashed lines mark the hit threshold in each screen. Relevant hits are labelled. **(E)** Gene Ontology Biological Process over-representation among significant resistance hits from the dBET6 screen with the V2 library, plotted as in Fig. 1D (matched Brunello screen). **(F)** sgRNA-level enrichment of the four recurrent indisulam resistance hits (ZMAT2, SETDB1, ZNF699, PPIH) in the Brunello (left) and V2 library (right) screens. Each vertical line is one sgRNA, positioned by its log2 fold change (indisulam vs DMSO); the greyscale ramp shows the binned density of all sgRNAs in the library. The dashed line indicates LFC=0. **(G)** sgRNA-level enrichment of the prominent UPS-associated resistance genes in the Brunello indisulam screen (DCAF15, C16ORF72, HUWE1, DDA1, GPS1, COPS8, GLMN), i.e. genes excluded from the V2 library. **(H)** Schematic of the competitive proliferation assay. **(I)** Competitive proliferation assay. Wildtype and knockout cells were mixed 1:1 and cultured with vehicle control (DMSO) or Indisulam (1 μM). All knockout lines and the corresponding AAVS1 control were cultured on the same plate and analysed in parallel for each timepoint; therefore, the same AAVS1 control curve is shown for all knockout comparisons. Competition assay data were analysed by 2-way repeated-measures ANOVA followed by Šídák’s multiple-comparisons test comparing each knockout with AAVS1 at each time point. *P < 0.05; **P < 0.01; ***P < 0.001; ****P < 0.0001. n = 2 biological replicates; each with 3 technical replicates per condition.

We next performed CRISPR screens using the V2 library across the full degrader panel described in Fig. 1B. Compared with the matched Brunello screens, the resistance landscape was markedly altered (Supplementary Fig. 1B). To determine whether *post hoc* removal of UPS-associated genes could similarly improve hit detection in conventional genome-wide screens, we excluded these genes from the Brunello dataset and repeated MAGeCK RRA analysis. Most significant resistance hits identified in the V2 screens failed to reach the same significance threshold in the corresponding Brunello screens. Indeed, significant V2 hits showed lower LFC values in the matched Brunello screens (Fig. 2C and Supplementary Fig. 1C). These results demonstrate that removing recurrent degrader-resistance genes at the library-design stage fundamentally reshapes the output of pooled TPD resistance screens and improves the detection of less penetrant resistance mechanisms, which cannot be simply recovered by excluding dominant UPS-associated hits during post hoc analysis.

Importantly, V2 screening uncovered resistance mechanisms with greater biological relevance to the degraded target (Fig. 2D and Supplementary Fig. 1D). For example, the THAL-SNS-032 (CDK9 degrader) V2 screen, but not the matched Brunello screen, identified INTS12, a subunit of the Integrator complex that antagonises CDK9-dependent transcriptional elongation ^18, 37^, as a significant resistance mechanism.

The move towards target related interpretable results is further evident in gene ontology analysis of significant resistance hits. Taking the dBET6 screens as an example, in which the top resistance hits were dominated by UPS-associated genes in Brunello screen (Fig. 1D), Gene Ontology analysis of the V2 screen revealed enrichment for biological processes linked to transcriptional regulation, consistent with the known role of BRD4 ^38, 39^ (Fig. 2E). We repeated the clonal dynamics analysis using experimentally derived growth values from the dBET6 screen (Supplementary Fig. 2A). Consistent with the computational modelling in Fig. 1A, the clonal dynamics analysis of the Brunello screen showed that UPS-associated resistance clones in which dBET6 activity was compromised, including CRBN, CAPRIN1, UBE2G1, GPS1, and COPS7B, rapidly expanded and compressed the relative abundance of the top V2 resistance hits. In contrast, removal of UPS-associated resistance genes in the V2 screen prevented the emergence of these dominant clones. Beyond PROTACs and molecular glue degraders, our library is compatible with tag-targeting protein degrader systems, including the widely used FKBP12^36V^ tag, as demonstrated by the identification of biological resistance mechanisms in CRISPR screens using cells expressing degron-tagged INTS12, a subunit of the Integrator complex^18^. This approach therefore provides a complementary screening framework for identifying dependencies that operate downstream of target degradation.

Across Brunello, V1, and V2 screens with the molecular glue degrader indisulam^10, 11^, a subset of resistance hits were consistently identified (Supplementary Fig. 2B). These included ZMAT2, PPIH, SETDB1, and ZNF699, which ranked as the top four resistance hits in the V2 screen. The recurrence of these genes across multiple screens nominated them as high-confidence mediators of indisulam resistance. Consistent with this, sgRNAs targeting each gene showed strong positive enrichment in indisulam-treated cells relative to DMSO controls in both the Brunello and V2 screens (Fig. 2F). In the Brunello screen, the enrichment of these genes was comparable to that of prominent UPS-associated resistance genes (Fig. 2G), demonstrating that under certain treatment conditions, strongly penetrant resistance mechanisms linked to target biology can emerge even in conventional genome-wide screens.

Nevertheless, the V2 library substantially improved the biological resolution of the indisulam screen. Gene ontology analysis of significant resistance hits from the Brunello screen was dominated by UPS-related processes, including protein deNEDDylation, ubiquitylation, and COP9 signalosome assembly, whereas the V2 screen revealed enrichment of additional biological pathways, including TORC1 signalling and autophagy (Supplementary Fig. 2C, D). Moreover, ZMAT2, despite showing resistance in the Brunello screen, became the top-ranked resistance hit in the V2 screen (Supplementary Fig. 2B) Together, these findings indicate that rational exclusion of UPS-associated resistance genes not only enables the detection of biological resistance mechanisms that may otherwise be masked, but also improves the prioritization and biological interpretability of target-related genetic interactions that are already detectable in conventional screens.

We next validated ZMAT2, PPIH, SETDB1, and ZNF699 as mediators of indisulam resistance using competitive proliferation assays (Fig. 2H, I). GFP positive cells nucleofected with sgRNAs targeting each candidate resistance gene were put in competition with tdTomato labelled control cells. This demonstrated that each of the resistance hits rapidly outcompeted the WT counterparts exclusively under indisulam treatment over the course of 10 days. AAVS1 sgRNA control cells remained in neutral competition with WT cells under indisulam treatment (Fig. 2I). In DMSO-treated conditions only SETDB1 loss conferred a competitive advantage, with all other hits demonstrating a modest decrease in growth (Supplementary Fig. 2E). This data validated that the loss of ZMAT2, PPIH and ZNF699 provides a selective advantage during indisulam treatment.

### ZMAT2 loss results in an attenuated transcriptional response to RBM39 degradation

Indisulam acts as a molecular-glue degrader that recruits the splicing factor RBM39 to the CRL4-DCAF15 E3 ligase for proteasomal degradation^10, 11^, resulting in widespread splicing defects and cytotoxicity. ZMAT2 and PPIH, are both components of the pre-catalytic spliceosomal B complex ^40, 41^ (Supplementary Fig. 3A), suggesting that modulation of spliceosomal activity downstream of RBM39 may offset its cytotoxic activity. To investigate the involvement of the spliceosome beyond these two factors we extracted all spliceosomal constituents from annotated subcomplexes using the CORUM database and intersected these with our V2 CRISPR screen hits. This uncovered that additional resistance hits were present in our indisulam screens linked to spliceosome function and splicing regulation, including RBM17 and PPIG (Fig. 3A). This data suggests that beyond the loss of ubiquitin proteasomal components, resistance to indisulam can arise from splicing-centred resistance mechanisms.

**Figure 3.**
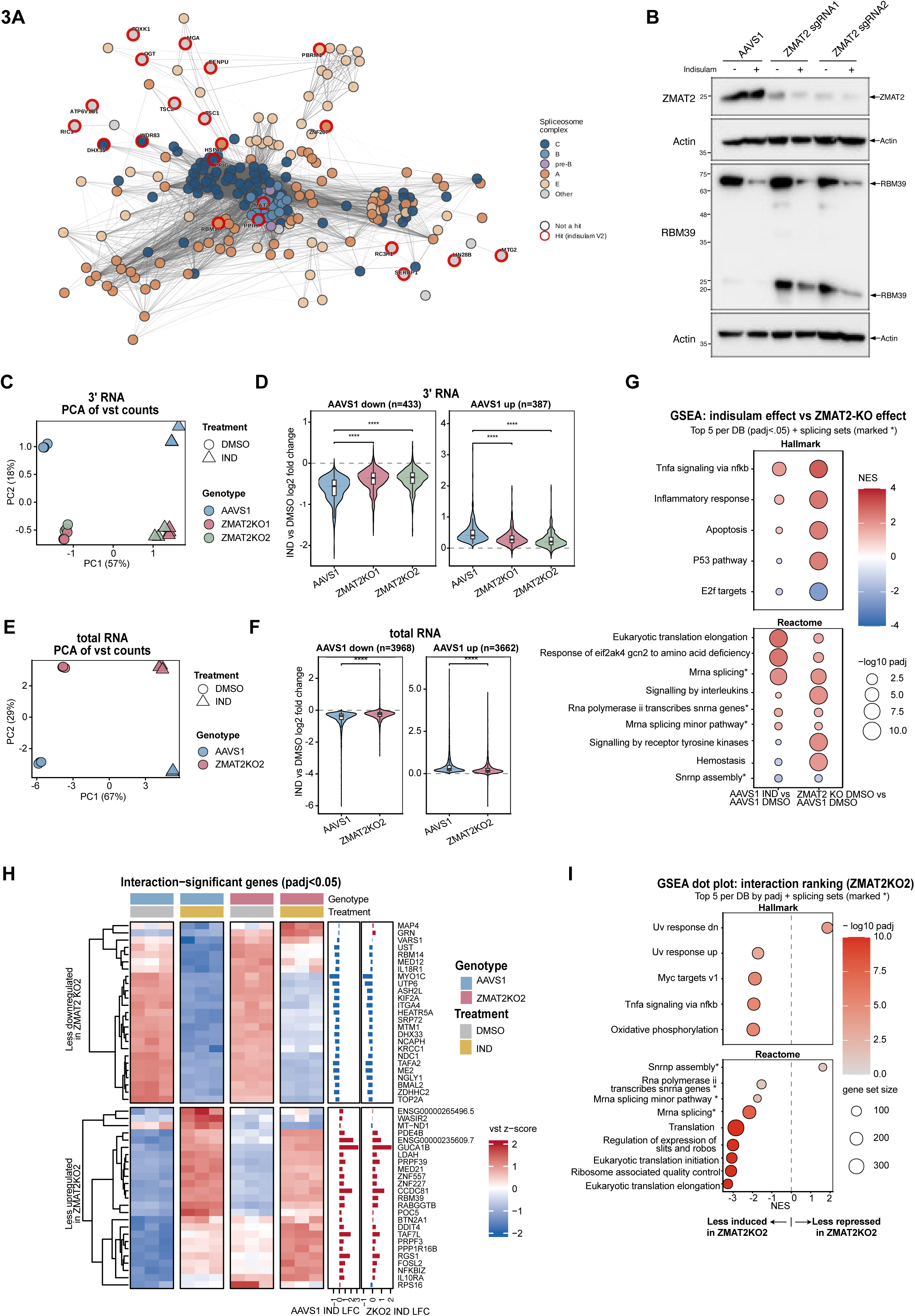
ZMAT2 loss blunts, but does not reverse, the indisulam-induced transcriptional response. (A) STRING interaction network of spliceosome proteins annotated by CORUM 4.0 and significant resistance hits from the V2 indisulam screen. Nodes are coloured by spliceosomal complex membership (E, A, pre-B, B, C or other). Membership defaults to the latest complex when dual membership is detected. Nodes outlined in red are significant resistance hits (p < 0.005, LFC > 1) in the V2 indisulam screen. (B) Western blot of ZMAT2 and RBM39 in KBM7 cells nucleofected with an AAVS1-targeting control sgRNA or two independent ZMAT2-targeting sgRNAs (sgRNA1 and sgRNA2) and treated with 1 µM indisulam or DMSO for 8h. (C) Principal component analysis of variance-stabilised 3′ RNA-seq from cells described in (B) (n = 3 replicates per condition), using the 500 most variable genes. (D) Violins plots of log2 fold change of significantly down-regulated (n = 433) or up-regulated (n = 387) in AAVS1 control cells, compared with ZMAT2 knockout cells (3′ RNA-seq). Indisulam-responsive genes were defined in AAVS1 cells (DESeq2, Benjamini–Hochberg adjusted-p < 0.05) and split into down-regulated (left) and up-regulated (right) sets. Paired Wilcoxon signed-rank test vs AAVS1; ****P < 0.0001, *P < 0.05 (E) Principal component analysis of variance-stabilised counts from deep total RNA-seq of KBM7 cells nucleofected with AAVS1 sgRNA control or sgRNA2 targeting ZMAT2, treated with 1 µM indisulam or DMSO for 8h. (n = 3 replicates per condition). (F) Violin plots of indisulam-induced log2 fold changes in genes significantly downregulated (n = 3,968) or upregulated (n = 3,662) in AAVS1 control cells, compared between AAVS1 control and ZMAT2-knockout cells (total RNA-seq). Indisulam-responsive genes were defined in AAVS1 cells (DESeq2, Benjamini–Hochberg adjusted-p < 0.05) and split into down-regulated (left) and up-regulated (right) sets. Paired Wilcoxon signed-rank test vs AAVS1; ****P < 0.0001, *P < 0.05. (G) Gene set enrichment analysis (GSEA) for total RNA-seq (fgsea, pre-ranked on the DESeq2 Wald statistic) of the indisulam response in AAVS1 cells (left column) and of the ZMAT2-knockout effect at baseline (ZMAT2KO2 vs AAVS1, DMSO; right column), across the MSigDB Hallmark and Reactome collections, with top 5 per database and splicing gene sets shown. Point colour, normalised enrichment score; point size, −log10(adjusted-p). (H) Heatmap showing row-wise z-scored variance-stabilised counts of top 25 (up and down) genes with significant genotype × treatment interactions in total RNA-seq (DESeq2, adjusted-p < 0.05), grouped by reduced or enhanced transcriptional responses to indisulam in ZMAT2-knockout cells. Bars indicate indisulam-induced log-fold changes (LFCs) in AAVS1 control and ZMAT2-knockout cells. (I) GSEA of the genotype treatment interaction (ZMAT2KO2), ranked on the interaction Wald statistic, across the two MSigDB collections, with top 5 per database and splicing gene sets shown; point colour, adjusted-p; point size, gene set size. Negative NES values indicate gene sets with attenuated upregulation in ZMAT2KO2 cells, whereas positive NES values indicate gene sets with attenuated downregulation. Dashed line, NES = 0.

Although splicing components were strongly enriched, it has been reported that disruption of splicing factors can impact the activity of molecular glue degraders and PROTACs via missplicing of E3 ligase components. For example, ILF3, a significant resistance hit in both the Brunello and V2 dBET6 screens, has been shown to reduce full-length CRBN expression by altering CRBN mRNA splicing and therefore acts upstream of target engagement ^42^, providing a rationale for excluding ILF3 from subsequent library versions. To examine whether ZMAT2 loss confers resistance by impairing indisulam-mediated RBM39 degradation, we performed Immunoblot analysis of RBM39 in the absence or presence of indisulam in AAVS1 or ZMAT2 knockout (KO) cells. This demonstrated that RBM39 degradation remained intact in ZMAT2 knockout cells following indisulam treatment (Fig. 3B). However, we did observe a short RBM39 isoform at around 25kDa, almost exclusively in the ZMAT2 knockout conditions. This shorter isoform was sensitive to indisulam-mediated degradation, indicating that it retains the indisulam binding region required for degradation. This isoform may arise from alternative splicing, altered TSS usage or termination or proteasomal cleavage as compared to full-length RBM39. Although this small isoform may act as a decoy for indisulam, we did not see a markedly reduced degradation of the full-length RBM39, suggesting that this is not the principal mechanism of resistance. Taken together, these data suggest that ZMAT2 loss confers resistance to the consequence of RBM39 degradation via compensation of its loss at the biological level possibly by altering spliceosomal and transcriptional activity directly.

We subsequently profiled the impact of ZMAT2 loss on the transcriptional response to indisulam by 3′ RNA-seq. Principal-component analysis (PCA) of the resulting data from ZMAT2-knockout and AAVS1-control cells treated with DMSO or indisulam revealed that samples separated primarily by treatment (PC1) and then by genotype (PC2) (Fig. 3C). In AAVS1-control cells, indisulam altered expression of 820 genes at adjusted-p < 0.05 and |LFC| > 0 (387 up, 433 down; DESeq2 Wald test) (Supplementary Fig 3B). Under vehicle conditions, ZMAT2 loss produced a comparatively modest and strongly up-skewed change in expression: 66 genes in KO1 (53 up, 13 down) and 61 genes in KO2 (55 up, 6 down) at adjusted-p < 0.05 and |LFC| > 0 (Supplementary Fig 3C). Upon indisulam treatment, ZMAT2 loss altered the expression of 396 genes in KO1 (189 up, 207 down) and 312 genes in KO2 (119 up, 193 down) at adjusted-p < 0.05 and |LFC| > 0 (Supplementary Fig 3B). To formally assess whether ZMAT2 loss modulates the transcriptional response to indisulam, we examined the genotype treatment interaction term. No genes reached significance in KO1 and only a single gene reached significance in KO2 (adjusted-p < 0.05). Interaction terms are often statistically under powered due to variance accumulation and the smaller effect sizes accompanied with assessing a difference of differences. Nevertheless, genes significantly induced or repressed by indisulam in AAVS1 cells displayed a compression of the corresponding IND-vs-DMSO fold change in both ZMAT2-KOs (Fig. 3D). The same trend was observed among all expressed genes that were significantly responsive to indisulam in both AAVS1 and ZMAT2KO genotypes (Supplementary Fig. 3D), indicating that the compression is unlikely due to regression to the mean, which may occur when defining the significant gene set based on a single comparison.

To capture the transcriptional response with greater sensitivity and resolution, we performed total RNA-seq analysis with ribodepletion and paired-end (150bp) sequencing. Analysis of the resulting data at the gene level resolved orthogonal treatment and genotype axes (Fig. 3E). Similar to the 3’RNA-seq data, the response to indisulam in AAVS1 cells was compressed in ZMAT2 knockout cells (Fig. 3F and Supplementary Fig. 3E). Assessing the effect of RBM39 loss and ZMAT2 loss alone, indisulam produced a large transcriptional response in control (AAVS1) cells (adjusted-p < 0.05, 3662 up, 3968 down; DESeq2 Wald test), and ZMAT2 KO alone perturbed a smaller set under DMSO conditions (adjusted-p < 0.05, 2259 up, 1996 down; DESeq2 Wald test) (Supplementary Fig 3F, G). Gene-set enrichment analysis of the indisulam effect in AAVS1 and of the ZMAT2 knockout in DMSO showed that RBM39 loss alone and ZMAT2 loss alone induces a coordinated stress and biosynthetic-remodelling program: translation and ribosome-machinery genes, the eIF2AK4/GCN2 amino-acid-deficiency (integrated stress) response, mRNA-splicing and snRNA-transcription sets, and inflammatory/TNFα-NF-κB signalling, while repressing E2F targets (proliferation) and the snRNP-assembly set (Fig. 3G). This is concordant with the same analysis in the 3′ RNA-seq (Supplementary Fig 3H), which showed that indisulam and ZMAT2 knockout produced a partially concordant baseline signature, likewise upregulating splicing, translation and inflammatory/TNFα sets and downregulating E2F targets and snRNP assembly (Fig. 3G and Supplementary Fig 3H), indicating that ZMAT2 loss may converge on the same mechanisms and cellular phenotypes as RBM39 loss.

Whilst in the 3′ RNA-seq, the genotype treatment interaction was essentially null, in the total RNA-seq the interaction identified 319 genes (219 up, 100 down) at adjusted-p < 0.05 (Fig. 3H). This is likely due to the deeper sequencing lending more statistical power. The interaction was mostly one of attenuation in both directions: genes normally repressed by indisulam were less repressed in the knockout, and genes normally induced by indisulam were less induced (Fig. 3H). This indicates that ZMAT2-knockout cells mount a blunted, not reversed, response to RBM39 degradation.

Gene-set enrichment on the interaction coefficient made this explicit and showed that the dominant, most significant signal was attenuated induction. The programs normally induced by indisulam (translation/ribosome genes, the integrated stress response, and mRNA-splicing/snRNA-transcription sets) were significantly less strongly induced in ZMAT2-knockout cells (negative interaction NES; Fig. 3I). Conversely, the snRNP-assembly set that indisulam normally represses was less strongly repressed (positive interaction NES).

Together, ZMAT2 knockout cells mount a muted transcriptional reaction to RBM39 loss, inducing the splicing-stress and translation-remodelling response to a lesser degree than AAVS1 control cells. Although indisulam still efficiently induced RBM39 degradation in ZMAT2-knockout cells, these cells responded to RBM39 loss in a blunted, non-reverting manner, from a baseline that already partly resembles the indisulam-perturbed state. These data place ZMAT2 loss downstream of RBM39 degradation and suggest that resistance may arise from rewired dependency on the splicing machinery rather than failure to engage the drug or reversal of its downstream program.

### ZMAT2 loss buffers indisulam-induced splicing defects

Our gene-level analyses (Fig. 3) showed that ZMAT2 knockout cells tolerate, rather than reverse, RBM39 degradation. Because indisulam is cytotoxic through inducing aberrant splicing, we next examined the drug response directly at this level using the total RNA-seq data. Principal-component analysis of exon-level counts separated samples primarily by treatment (PC2) and genotype (PC1), as the gene level analysis. Notably, indisulam-treated ZMAT2 knockout cells sat closer to DMSO-treated samples than their indisulam-treated AAVS1 counterparts (Fig. 4A), suggesting ZMAT2 loss partially attenuates the splicing response to RBM39 degradation.

**Figure 4.**
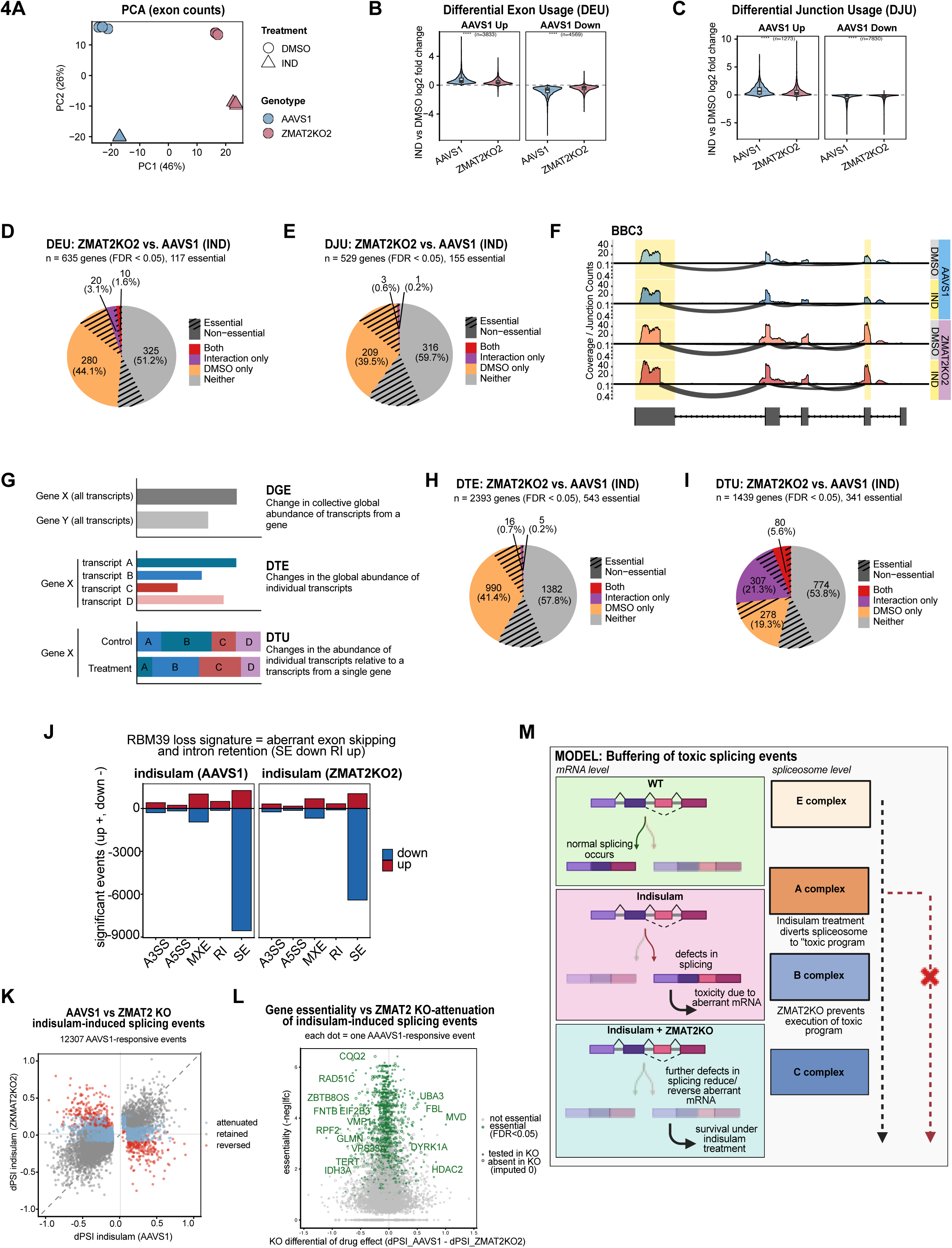
ZMAT2 loss buffers indisulam-induced splicing defects. **(A)** Principal component analysis of total RNA-seq log-CPM exon counts (top 500 most variable exons) for AAVS1 and ZMAT2KO2 cells treated with indisulam or DMSO (3 replicates per condition) **(B)** Violin plots of significant differential exon usage (DEU) in the AAVS1 treatment response (edgeR, adjusted-p < 0.05) shown in the two genotypes, split by up and down log-fold change directions; ****P < 0.0001 **(C)** Violin plots of significant differential junction usage (DJU) events in the AAVS1 treatment response (ASpli, adjusted-p < 0.05) shown in the two genotypes, split by up and down log-fold change directions; ****P < 0.0001 **(D)** Classification of DEU significant genes (edgeR, Simes test for gene-level significance, adjusted-p < 0.05) in the ZMAT2KO2 vs. AAVS1 under treatment contrast. Genes are classified as “Interaction only” or “Baseline (DMSO only)” if it is also significant in the interaction contrast, or the ZMAT2KO2 vs. AAVS1 under DMSO contrast (but not both), or “Both” when it is significant in all three contrasts. The remaining genes that show significance between ZMAT2KO2 and AAVS1 under indisulam but do not fall under these three categories are either false discoveries or those that fail to reach statistical significance due to lack of statistical power inherent to these models. Proportions of essential genes were further illustrated with dashed line covered area. Here essential genes are defined from Blomen et al., 2015 ^43^, which defines essentiality in KBM7 cells. **(E)** As in **(D),** for differential junction usage significant genes reported by ASpli (adjusted-p < 0.05) **(F)** Sashimi plot of gene BBC3, with merged exons significant in the genotype effect under DMSO highlighted in yellow. Total RNA-seq read coverage across the gene locus is shown for both genotype and treatment condition. Each row is the mean RPKM-normalised coverage of the group. Below the zero line, arcs indicates junction reads, with arc height and line-width both scaling to the junction’s RPM-normalised read count summed across replicates. The collapsed gene model with overlapping exons merged is shown at the bottom. **(G)** Schematic of the differential gene expression (DGE) / differential transcript expression (DTE) / differential transcript usage (DTU) framework. **(H)** As in **(D)**, for DTE (edgeR, adjusted-p < 0.05) **(I)** As in **(D)**, for DTU (edgeR, adjusted-p < 0.05) **(J)** Composition of significant alternative splicing events by class and direction (rMATS-turbo). Bars show the number of significant events (FDR < 0.05, |ΔPSI| > 0.1, JC) for each of the five event classes (A3SS, A5SS, MXE, RI, SE) in two contrasts: indisulam in AAVS1 and indisulam in ZMAT2KO2. Red, events with more inclusion or retention; blue, fewer. A3SS, alternative 3’ splice site; A5SS, alternative 5’ splice site; MXE, mutually exclusive exons; RI, retained intron; SE, skipped exon. **(K)** Event-level attenuation of the indisulam splicing response by ZMAT2 loss. For the 12,307 splicing events significantly responsive to indisulam in AAVS1 cells, ΔPSI in ZMAT2KO2 cells is plotted against ΔPSI in AAVS1 cells. Events are classified as, retained (significant (FDR < 0.05, |ΔPSI| > 0.1) in both and same sign) or reversed (significant in both opposite sign) and attenuated (significant only in AAVS1 indisulam comparison. PSI: Percent Spliced In **(L)** CRISPR screen derived essentiality of genes containing indisulam-induced splicing events, plotted against the attenuation of that event by ZMAT2 loss (ΔPSI_AAVS1 − ΔPSI_ZMAT2KO2). Each pointis one AAVS1-significant event. Green points, essential genes (FDR < 0.05 in the CRISPR fitness screen – DMSO vs T0); open circles, events absent in the knockout (imputed as 0). **(M)** Model. RBM39 degradation by indisulam generates aberrant transcripts, a subset of which are toxic. ZMAT2 loss does not abolish RBM39 degradation but reduces aberrant products, buffering the aberrant splicing and transcriptional consequences of RBM39 loss.

To quantify the impact of ZMAT2 loss on the splicing response to indisulam, we assessed differential exon usage (DEU; edgeR) and differential junction usage (DJU; ASpli) (Fig. 4B, C). The exons and junctions significantly responsive to indisulam in the AAVS1 cells exhibited attenuated treatment-induced changes in ZMAT2 knockout cells, with reduced magnitude observed for both increased and decreased exon/junction usage, mirroring the compression observed at the gene expression level. Consistent with this, the compression of exon usage in ZMAT2 knockout cells was also observed when analysing treatment-effect log-fold change for all expressed exons, with the majority of exons significantly responsive to indisulam in both genotypes lying between the identity line (y = x) and the x-axis (Supplementary Fig. 4A).

We further classified genes that showed differential exon usage between ZMAT2 knockout cells and AAVS1 cells upon indisulam treatment by whether they are also significant under DMSO condition (i.e. ZMAT2 knockout versus AAVS1 under DMSO, indicating a baseline genotype effect), and whether they are significant in the interaction contrast. As shown in Fig. 4D, 290 genes are significant in the DMSO condition, which includes 10 genes that are also significant in the interaction contrast (280 “DMSO only” and 10 “both” in the pie chart), and 30 genes (20 “interaction only” and 10 “both”) were also significant in the interaction contrast. As illustrated in Supplementary Fig 4B, significant events in the ZMAT2 knockout cells vs. AAVS1 cells upon indisulam treatment, unless are false positives, can be attributed as either knockout effect present under DMSO or interaction effect, or a combination of both. Yet, in our case, over half of the genes (325, 51.2%) cannot be explained, as with the 3′ RNA-seq analysis, this likely reflects that interaction terms are underpowered by variance accumulation. Cross-referencing with essentiality data from Blomen et al., 2015^43^ revealed essential genes within each category (shaded sectors in the pie chart), suggesting candidate genes through which altered exon usage may contribute to indisulam resistance conferred by ZMAT2 loss. Likewise, among the 529 genes with differential junction usage between ZMAT2 knockout vs AAVS1 knockout cells under indisulam treatment, we identified 4 gene (3 “interaction only” and 1 “both”) that showed a significant genotype × treatment interaction (Fig. 4E).

Consistent with its role as a human spliceosome B component ^40, 41^, the genotype effect of ZMAT2 is evidenced by the alternative splicing of the pro-apoptotic gene BBC3 (PUMA), which showed differential usage of two exons attributable to ZMAT2 knockout at baseline (DMSO; Fig. 4F). ZMAT2 loss also altered the splicing of RBM39 itself. RBM39 exons showed significant treatment, genotype, and genotype × treatment interaction effects (Supplementary Fig. 4C, D). A single annotated shorter RBM39 transcript isoform, RBM39-237, was significantly upregulated in ZMAT2-knockout cells (Supplementary Fig. 4E).

We next performed two transcript-level analyses of differential transcript expression (DTE) and differential transcript usage (DTU) (Fig. 4G-I). Under indisulam treatment, 2393 genes showed differential transcript expression between ZMAT2 knockout cells and AAVS1 cells (Fig. 4H), of which 21 genes (16 “interaction only” and 5 “both”) were significant under the genotype × treatment interaction model (Fig. 4H). Differential transcript usage displayed a distinct landscape, with 387 (307 “interaction only” and 80 “both”) out of 1,439 genes showed significant genotype × treatment interaction effects (Fig. 4I). Since DTU quantifies the relative abundance of transcript isoforms within a gene (Fig. 4G), it removes the large indisulam-driven changes in total gene expression levels improving statistical detection. Ultimately, these findings suggest that under indisulam treatment, ZMAT2 loss reshapes transcript usage and reduces production of specific aberrant transcripts induced by RBM39 loss.

To further classify the effect on splicing alterations independently of annotated transcript isoforms, we tested the treatment response in the two genotypes individually using rMATS-turbo^44^, which does not support interaction/complex model designs (Fig. 4J). In AAVS1 cells, indisulam reproduced the established RBM39-loss signature: a transcriptome-wide shift toward cassette-exon exclusion (lower percent-spliced-in; SE-down; exon skipping) and increased intron retention (RI-up), with comparatively few changes in the mutually-exclusive-exon and alternative-5′/3′-splice-site classes (Fig. 4J). Both signatures were markedly blunted in ZMAT2-knockout cells: the number of significant events fell in every class and most steeply for exon skipping, indicating that ZMAT2 loss reduces the number of exons indisulam drives to exclusion rather than redirecting them. At the event level (Fig. 4K), the majority of the 12,307 indisulam-responsive changes in AAVS1 cells retained their direction in ZMAT2-knockout cells but at reduced magnitude, with a defined subset collapsed toward zero (attenuated/lost) and a smaller subset changed sign (reversed). Because the bulk program is directionally preserved, this buffering is selective rather than a global loss of splicing, consistent with ZMAT2 loss lowering the yield of specific indisulam-induced events.

To determine whether these buffered splicing events in ZMAT2 knockout cells could plausibly account for indisulam resistance, we intersected the indisulam-induced splicing events most strongly attenuated by ZMAT2 loss with gene essentiality in KBM7 cells. Whereas our previous analyses used the essentiality data from Blomen et al., 2015^43^, here we instead used the DMSO vs T0 comparison from our CRISPR screen to derive gene essentiality scores. This provides quantitative fitness estimates measured in the same experimental context as the drug screens, providing better rankings for candidate splicing events affecting survival. A subset occurred in essential genes (Fig. 4L), nominating candidate splicing events whose attenuation could contribute to indisulam resistance.

Based on these findings we proposed a model (Fig. 4M) in which indisulam-induced RBM39 degradation causes splicing defects and promotes the generation of aberrant transcripts, a subset of which are toxic. Loss of the core human spliceosome B complex component ZMAT2 does not prevent RBM39 degradation, rather attenuates these aberrant splicing events, thereby reducing these toxic transcripts and buffering the consequences of RBM39 loss.

## Discussion

Targeted protein degradation (TPD) has emerged as a promising therapeutic and research strategy by enabling rapid, selective and reversible protein depletion. However, its application in genetic screenings has been limited because disruption of the ubiquitin-proteasome system (UPS) confers dominant resistance by preventing target degradation, thereby masking bona fide biological genetic interactions ^12, 27^. Here, we developed a TPD-compatible CRISPR library that selectively removes UPS-mediated degrader-resistance genes while retaining near-genome-scale coverage. We demonstrate that this strategy enables the identification of biological resistance mechanisms acting downstream of target degradation across multiple TPD compounds. As proof of principle, we identified the spliceosome B component ZMAT2 as a major resistance mechanism to RBM39 degradation by the molecular glue degrader indisulam, whereby ZMAT2 depletion buffers the downstream splicing defects induced by RBM39 loss. Together, our work establishes a generalizable framework for separating resistance to the degradation process from resistance to the biological consequences of target depletion.

The predominance of UPS components in TPD-based CRISPR screens can be explained by the clonal dynamics of pooled genetic screens. Our *in-silico* simulations show that, within a mixed population of resistant clones, loss of the UPS machinery confers the greatest growth advantage because the targeted protein degradation is abolished entirely. These degrader-resistance clones rapidly outcompete biologically resistant clones with less penetrant phenotypes, ultimately dominating the screened population. Because CRISPR screen readouts are inherently compositional, expansion of degrader-resistance clones proportionally reduces the abundance of other resistance clones. Therefore, simply removing predictable UPS hits during downstream bioinformatics analysis is not sufficient, as these biological-resistance clones never reach detectable abundance. Instead, depleting UPS-mediated degrader-resistance genes from the screening library is required to reveal the underlying target-dependent genetic interactions.

The UPS-related genes removed from our TPD-compatible library were compiled from published literature, our own screening data, and manual curation of E3 ligases and their associated regulatory components. As E3 ligases and their associated regulatory networks continue to expand ^45^, the exclusion list should be iteratively updated to incorporate newly identified components of the TPD machinery. In turn, recurrent resistance hits identified from TPD-based CRISPR screens may themselves uncover previously unrecognized components or regulators of the UPS, thus expanding the UPS network. It will be important to validate some of the nominated degrader failure candidates such as UBE2K, PEX10 and NSFL1C. As new TPD compounds and tag-based degradation systems emerge, the excluded gene set can be tailored to compounds of interest. For example, tag-based degrader screens may benefit from excluding factors such as FKBP1A, which influence degradation systems based on FKBP-derived tags or ligands. Likewise, analogous libraries could be generated for lysosome-based degraders^1^ by excluding genes involved in lysosomal degradation pathways.

Although developed for TPD-based CRISPR knockout screens, our TPD-compatible library could be applied to other functional genomic approaches such as CRISPR activation (CRISPRa) and CRISPR interference (CRISPRi), when combined with TPD compounds. More broadly, our study illustrates a conceptual strategy for improving genetic screens by removing dominant, predictable determinants of compound activity, thereby unmasking biological genetic interactions. This principle could be extended to other drug-based CRISPR screens by excluding genes required for drug efflux pumps and enzymatic activation steps required for drug action (e.g. AraC resistance^46^). More generally, the same concept may also improve signalling-based genetic screens. For example, IFNg-driven PD-L1 FACS screens are typically dominated by loss-of-function mutations in core IFNg pathway components such as PD-L1, IFNGR1 and STAT1 ^47, 48^, which can obscure additional regulators with more subtle phenotypes. Designing pathway-focused libraries that exclude well-established dominant regulators may therefore provide a general strategy for uncovering novel biological mechanisms across diverse screening platforms.

Our data showed that the TPD-compatible library, compared with the conventional genome-wide CRISPR Brunello library, substantially enriches for biological rather than UPS-related resistance hits across multiple TPD compounds. Notably, the top four indisulam resistance hits, ZMAT2, PPIH, SETDB1 and ZNF699, were also significantly enriched in the Brunello screen, albeit to a lesser extent. This suggests that highly penetrant biological resistance mechanisms can still be detected in conventional genome-wide screen settings. Nevertheless, the TPD-compatible library markedly improves their prominence and interpretability by reducing the number of competing degrader-failure mechanisms.

We validated ZMAT2 as a top resistance mechanism to indisulam treatment. Indisulam acts as a molecular glue by recruiting RBM39 to DCAF15, resulting in RBM39 ubiquitination, proteasomal degradation and splicing defects ^10, 11^. Importantly, RBM39 degradation remained intact in ZMAT2-knockout cells, excluding the possibility that ZMAT2 functions as a UPS component or a regulator of UPS proteins that is required for degrader activity. ZMAT2 is a component of the human B spliceosome complex ^40^, and notably, many of the top resistance hits identified in the indisulam screen were also spliceosome-associated proteins (Fig. 3A), suggesting that altered spliceosome composition or progression through spliceosome assembly modulates cellular sensitivity to RBM39 loss. ZMAT2 deficiency broadly attenuated the transcriptional response to indisulam. Differential exon usage and differential junction usage also demonstrated a global attenuation of indisulam-induced splicing defects. Splicing analysis further demonstrated the loss and or reversal of a subset of indisulam induced defects in the presence of ZMAT2 knockout, favouring a model where ZMAT2 knockout directly lowers the yield of toxic indisulam induced splicing defects, allowing cells to tolerate indisulam treatment. We also observed a short RBM39 isoform in ZMAT2-knockout cells, irrespective of indisulam treatment. This isoform remained susceptible to indisulam-induced degradation, suggesting that it retains the structural features required for DCAF15-mediated recruitment. The origin and functional significance of this short RBM39 isoform remain unclear. Determining whether this isoform retains spliceosomal activity and contributes to the resistant phenotype will be an important direction for future investigation.

In summary, we developed a TPD-compatible CRISPR library that enables the discovery of biological resistance mechanisms that are otherwise masked by degrader-failure mechanisms. More broadly, our work establishes a general framework for rational library design in functional genomics, demonstrating that removing dominant, predictable determinants of compound activity can reveal weaker but biological genetic interactions. We anticipate that this principle will be broadly applicable across diverse targeted protein degradation modalities, chemical perturbations and genetic screening platforms.

## Figure legends

**Supplementary Figure 1.**
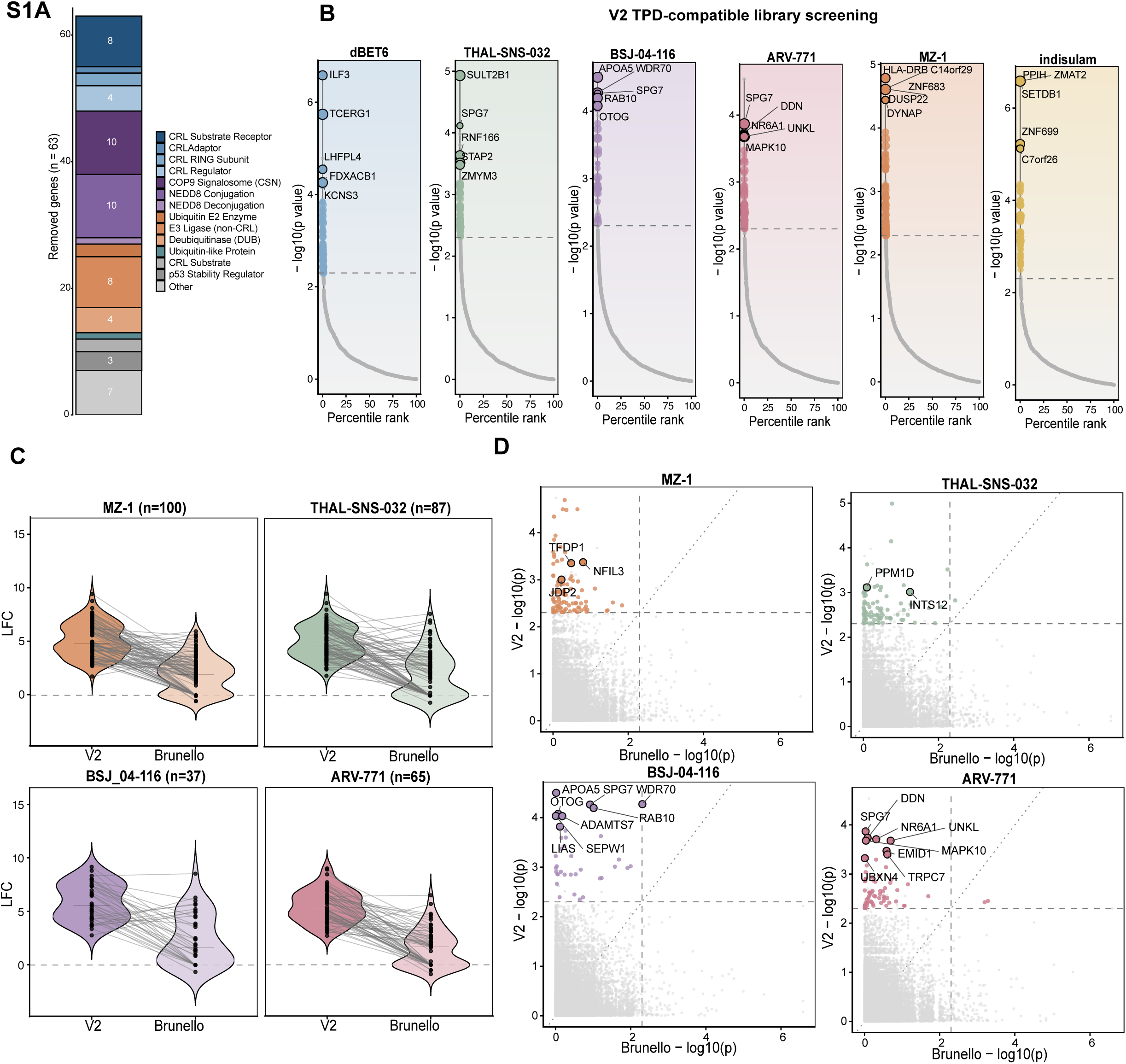
Design and composition of the TPD-compatible libraries. **(A)** Functional classification of the 63 genes excluded from the V1 library, by UPS category. **(B)** Ranked significance plots of the six V2 library resistance screens, as in Fig. 1C. **(C)** Gene-level log2 fold change (drug vs DMSO) of significant V2 resistance for MZ-1, THAL-SNS-032, BSJ-04-116 and ARV-771, plotted as in Fig. 2C. **(D)** Significance concordance between Brunello and V2 screens for MZ-1, THAL-SNS-032, BSJ-04-116 and ARV-771, plotted as in Fig. 2D.

**Supplementary Figure 2.**
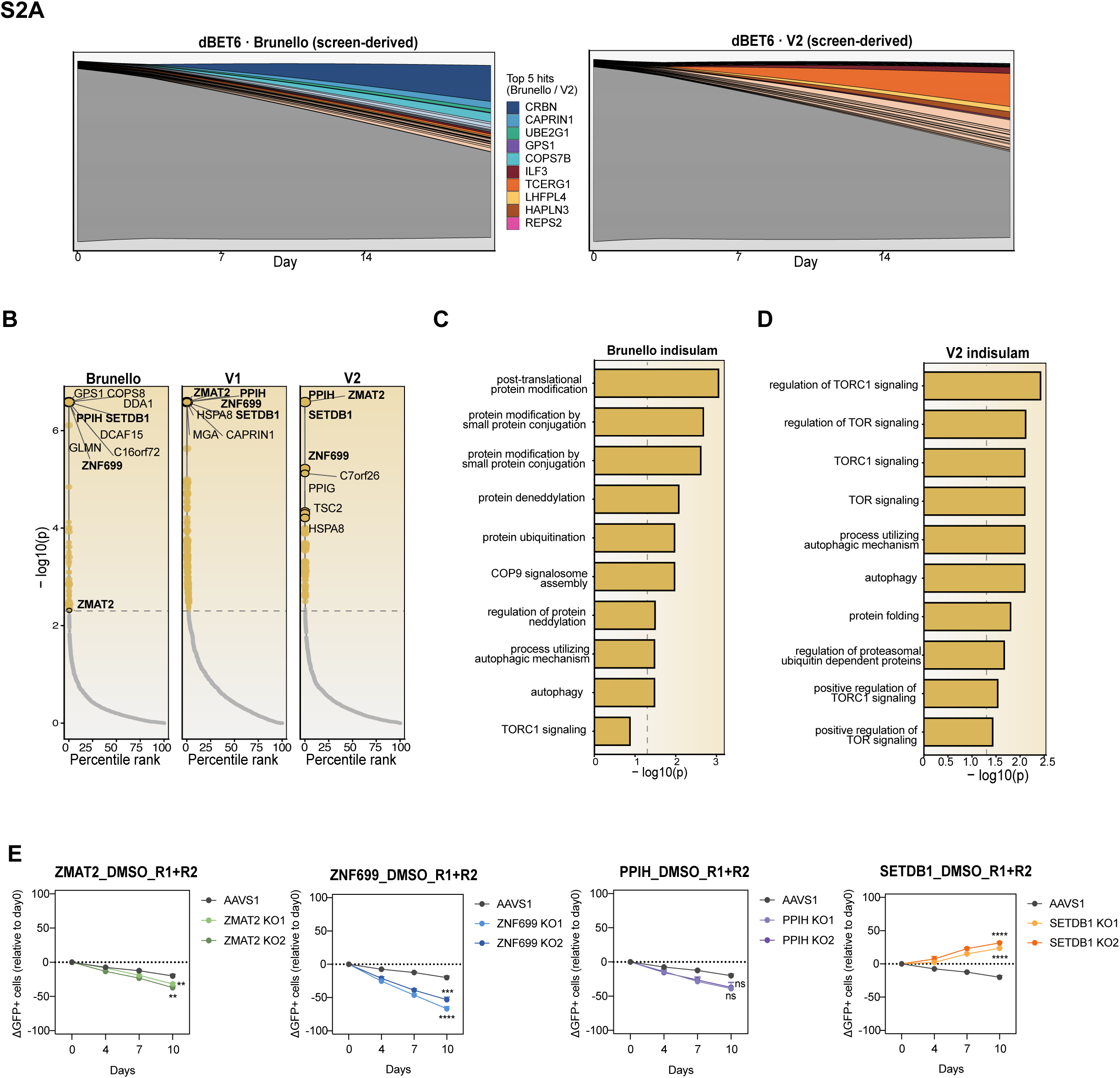
Screen-derived clonal dynamics and the recurrent indisulam resistance hits. **(A)** Clonal dynamics simulated from experimentally derived growth values from the dBET6 screens. Each gene’s MAGeCK positive-selection log2 fold change was converted to a per-day growth advantage over background and clones propagated deterministically over 21 days. Left, Brunello screen; right, V2 screen. The top five clones per class by MAGeCK score are individually shown. **(B)** Ranked significance plots of the indisulam screens with the Brunello, V1 and V2 libraries. ZMAT2, PPIH, SETDB1 and ZNF699 are labelled in each screen. **(C, D)** Gene Ontology Biological Process over-representation among significant indisulam resistance hits in the Brunello (C) and V2 (D) screens. **(E)** Competitive proliferation assay under vehicle (DMSO), as in Fig. 2I. All knockout lines and the corresponding AAVS1 control were cultured on the same plate and analysed in parallel for each timepoint; therefore, the same AAVS1 control curve is shown for all knockout comparisons. Competition assay data were analysed by 2-way repeated-measures ANOVA followed by Šídák’s multiple-comparisons test comparing each knockout with AAVS1 at each time point. *P < 0.05; **P < 0.01; ***P < 0.001; ****P < 0.0001. n = 2 biological replicates; each with 3 technical replicates per condition.

**Supplementary Figure 3.**
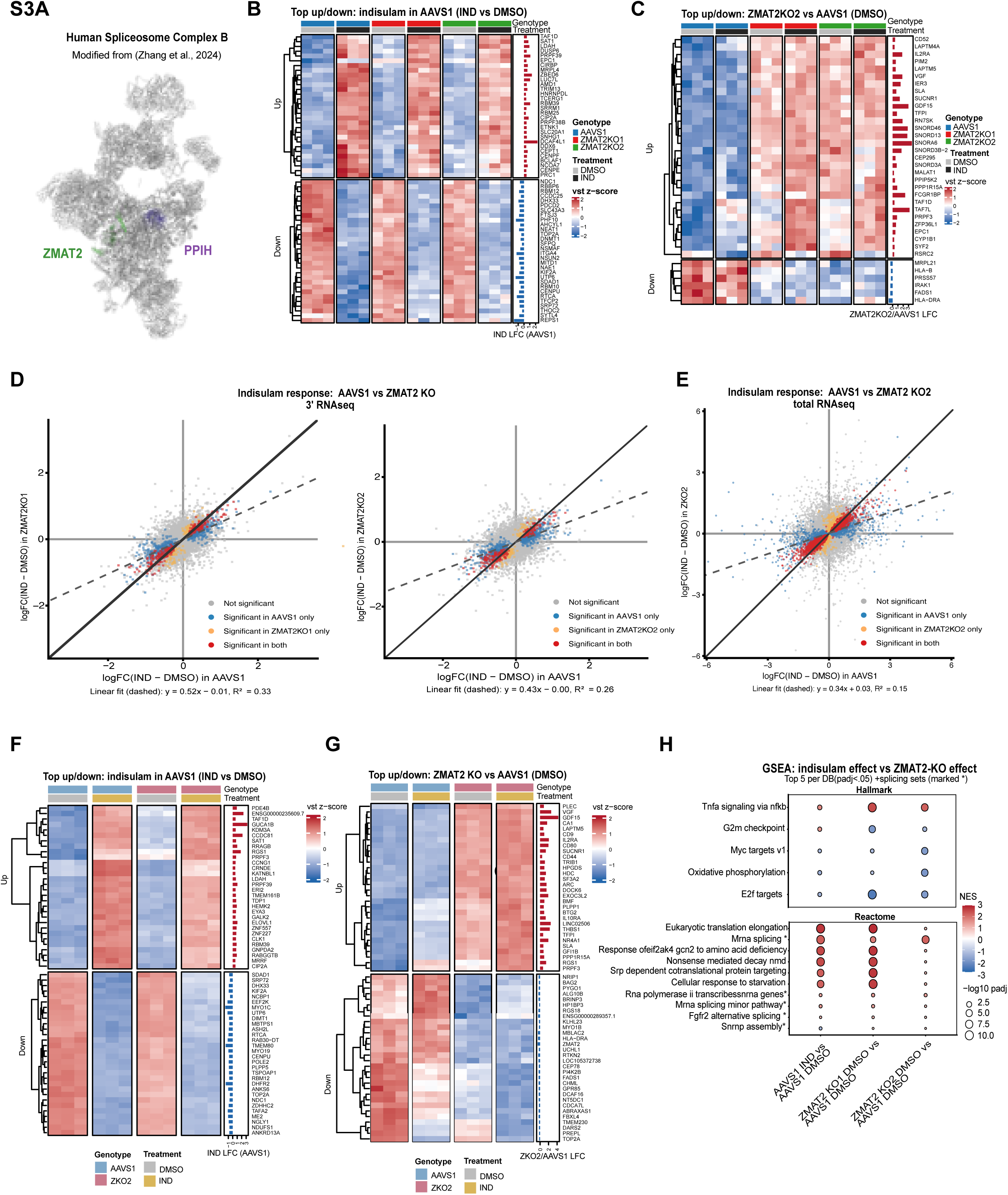
Transcriptional consequences of indisulam treatment and ZMAT2 loss. **(A)** ZMAT2 (green) and PPIH (purple) are components of the human spliceosome B complex. Structure modified from Zhang et al., 2024^40^. **(B)** Top 30 up– and down differentially expressed genes following indisulam treatment compared with DMSO in AAVS1 cells (3′ RNA-seq; DESeq2, adjusted-p < 0.05). Heatmap shows row-wise z-scored variance-stabilised counts across all three genotypes and both treatments. Right, the indisulam-vs-DMSO log2 fold change in AAVS1. **(C)** Up to top 30 up– and down differentially expressed genes between ZMAT2KOs and AAVS1 in DMSO condition (adjusted-p < 0.05; DESeq2 Wald test). Heatmap shows row-wise z-scored variance-stabilised counts across all three genotypes and treatments. Right, the ZMAT2KO2-vs-AAVS1 log2 fold change in DMSO. **(D)** Scatter plots comparing the differential gene expression (3′ RNA-seq) to indisulam between AAVS1 and ZMAT2KO cells. All expressed gene level LFCs (indisulam vs DMSO) in AAVS1 cells (x-axis) and ZMAT2KO clone cells (y-axis) are plotted. Points are coloured by differential expression at padj < 0.05 (Benjamini–Hochberg) within each genotype: significant in AAVS1 treatment effect only (blue), ZMAT2 KO treatment effect only (orange), both (red), or neither (grey). The solid line is y = x. Compression of the indisulam response is observed with most genes differentially expressed under indisulam in both genotypes (red) falling between y=x and the x axis. The dashed line is an ordinary least-squares fit of all expressed genes. **(E)** As in **(D).** Scatter plots comparing the differential gene expression (total RNA-seq) to indisulam between AAVS1 and ZMAT2KO2 cells. All expressed gene level LFCs (indisulam vs DMSO) in AAVS1 cells (x-axis) and ZMAT2KO2 clone cells (y-axis) are plotted. **(F, G)** As in **(B, C),** for the total RNA-seq dataset (AAVS1 and ZMAT2KO2). **(H)** GSEA of the indisulam response in AAVS1 and of the baseline ZMAT2-knockout effect in each of the two knockout clones (3′ RNA-seq), across the Hallmark and Reactome collections. Plotted as in Fig. 3G.

**Supplementary Figure 4.**
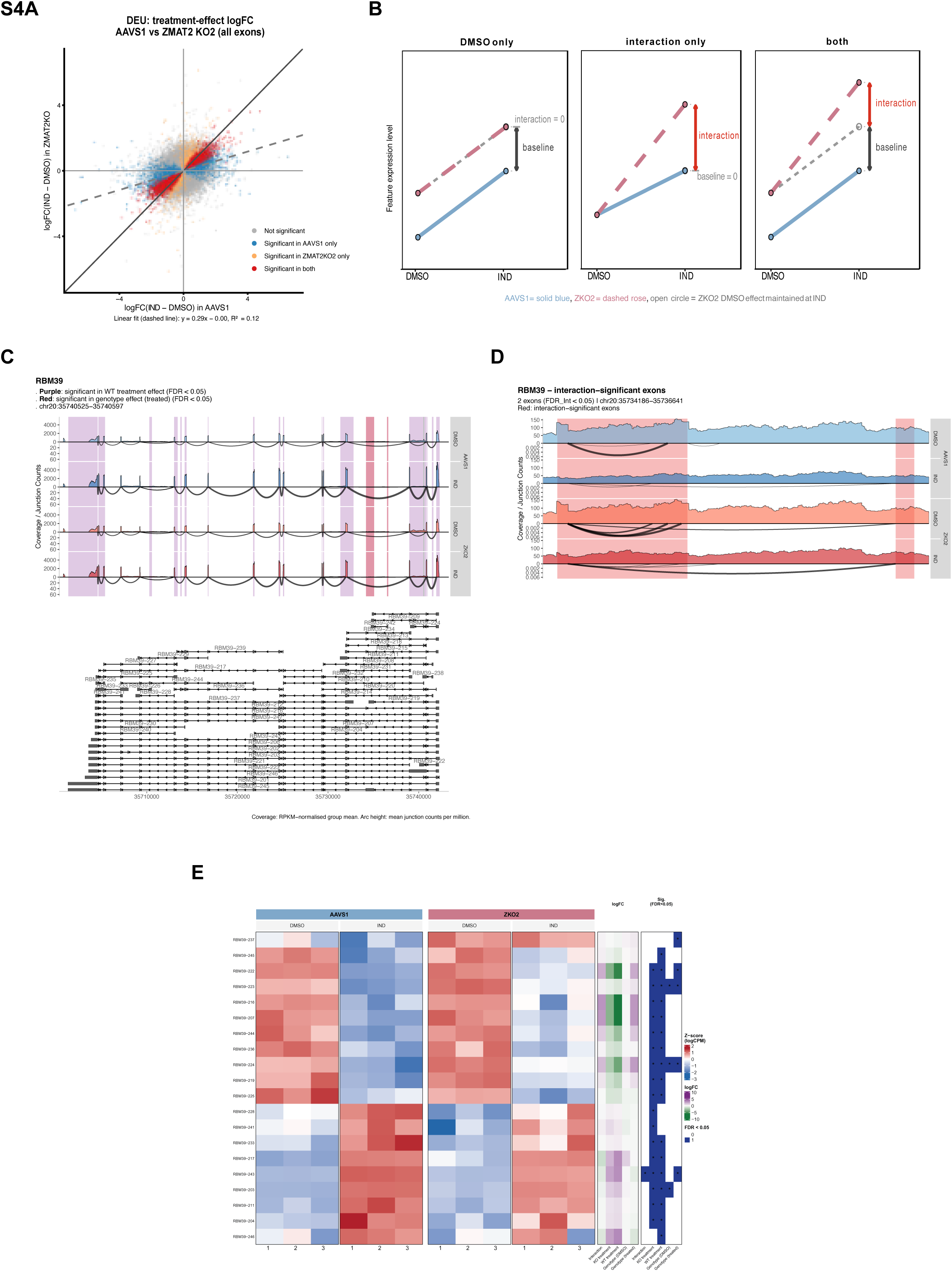
ZMAT2 loss alters RBM39 transcript isoform usage. **(A)** Scatter plot comparing the differential exon usage to indisulam between AAVS1 and ZMAT2KO2 cells. All exon level LFCs (indisulam vs DMSO) in AAVS1 cells (x-axis) and ZMAT2KO2 cells (y-axis) are plotted. Points are coloured by differential exon usage at padj < 0.05 (Benjamini–Hochberg) within each genotype: significant in AAVS1 treatment effect only (blue), ZMAT2 KO2 treatment effect only (orange), both (red), or neither (grey). The solid line is y = x. Compression of the indisulam response is observed with most exons differentially used under indisulam in both genotypes (red) falling between y=x and the x axis. The dashed line is an ordinary least-squares fit of all exons tested. **(B)** Illustration of all three possible cases for a significant feature in the ZMAT2KO2 vs. AAVS1 under indisulam comparison in an ideal setting with no false discoveries and sufficient statistical power. The effect found under treatment can be attributed to the knock-out effect that were also present under DMSO (“DMSO only”), or an interaction effect between genotype and treatment (“interaction only”), or a combination of both. The pie charts in Fig. 4 categorize genes into these scenarios by checking significance across comparisons, and the genes not in any of these cases (in grey) are either false discoveries (in the ZMAT2KO2 vs. AAVS1 under indisulam comparison) or genes that fail to reach statistical significance. It should be noted that the interaction term has lower statistical power inherently, which likely caused genes with interaction effect failing to reach significance. **(C)** Sashimi plot of gene RBM39, with significant merged exons highlighted. Total RNA-seq read coverage across the gene locus is shown for both genotype and treatment condition. Each row is the mean RPKM-normalised coverage of the group. Below the zero line, arcs indicate junction reads, with arc height and linewidth both scaling to the junction’s RPM-normalised read count summed across replicates. The gene model is shown at the bottom. The purple highlighting indicates merged exons significant in the AAVS1 treatment effects. The red highlighting indicates the merged exons significant in the genotype effect of the treated samples. The two merged exons in red are also the only two exons significant in the interaction contrast, and both are significant in the AAVS1 treatment effects as well. A zoomed in view of these exons are shown in (D). **(D)** Zoomed-in view of the two exons highlighted in red in (C), which are significant in the genotype effect of treated samples, the interaction contrast and the AAVS1 treatment effects. **(E)** Heatmap of RBM39 transcript relative expression (z-score of logCPM) across replicates and the log fold change in each contrast.

## Materials and Methods

### Cell lines and culture conditions

KBM7 cells were cultured at 37°C and 5% carbon dioxide (CO2) in IMDM supplemented with 10% fetal bovine serum (FBS), 100U/mL penicillin, 100 μg/mL streptomycin. Cells were authenticated using short-tandem-repeat (STR) profiling and were also tested for mycoplasma detection (Peter McCallum Cancer Centre service).

### V1 and V2 library synthesis and cloning

The sgRNA spacer sequences from the Brunello Human CRISPR knockout pooled library (Addgene #73179) were used as the starting point for library design. To generate the TPD-compatible V1 library, sgRNAs targeting the 63 genes excluded as described in the main text were manually removed, resulting in a library containing 77,189 sgRNA spacer sequences. To generate the TPD-compatible V2 library, sgRNAs targeting an additional 40 genes identified through iterative library refinement and described in the main text were subsequently removed, resulting in a final library of 77,022 sgRNA spacer sequences targeting 19,011 genes. The corresponding oligonucleotide pools were synthesised (GenScript), PCR-amplified, and cloned into the LentiGuide-Hygro vector (Addgene #139462) using BsmBI (NEB) restriction sites. Complete removed gene lists are provided in Supplementary Table 1 (V1) and Supplementary Table 2 (V2) and sgRNA sequences for V2 are provided in Supplementary Table 3 (V3).

### CRISPR-Cas9 screens

KBM7 cells stably expressing humanized S.pyogenes Cas9 endonuclease were generated by lentiviral transduction with the FU-Cas9-Cherry vector (Addgene #70182), followed by FACS-selection for mCherry-positive cells. Lentiviral for the Brunello sgRNA library (Addgene #73178) and the TPD-compatible CRISPR knockout libraries (V1 and V2; generated in-house) were produced using polyethylenimine (PEI; Sigma-Aldrich) transfection reagent. KBM7 cells expressing Cas9-Cherry were transduced with library virus at a optimal multiplicity of infection (MOI) of 0.3 and a fold representation of 250 for each individual sgRNA. Transduced cells were selected with 1 μg/mL puromycin for 7 days, after which T0 baseline pellets were collected, and the remaining population was split into relevant treatment conditions (DMSO, Indisulam 500nM, ARV771 40nM, MZ1 200nM, dBET6 20nM, THAL-SNS-032 150nM, BSJ-04-116 40nM) for a total of 21 days and cell pellets were collected as Tend samples. Genomic DNA was extracted using the QIAamp DNA Blood Midi Kit (Qiagen, Cat. #51185). PCR amplification of sgRNA sequences from genomic DNA was performed using Ex Taq DNA Polymerase (Takara, Cat. #RR001C) according to the Broad Institute protocol. Amplified libraries were sequenced at the WEHI Genomics Facility on an Illumina NextSeq 2000 using single-end 100-cycle sequencing.

### Competition Proliferation Assays

KBM7 cells stably expressing either eGFP or tdTomato fluorescent reporters introduced by PiggyBac transposition (PB-CAG-eGFP, Addgene #40973; PB-CAG-tdTomato, Addgene #133569) were nucleofected with Alt-R S.p. HiFi Cas9 Nuclease V3 and sgRNAs targeting AAVS1(spacer sequence 5’->3’: GTCACCAATCCTGTCCCTAG), ZMAT2 (spacer sequence 5’->3’ ZMAT2KO1: TCTCGCTTGACAGGCTGCAC; ZMAT2KO2: CGTAAAAGCTCTCGCTTGAC), PPIH (spacer sequence 5’->3’ PPIH KO1: TCTGCTCAGGGAGATGGTAC; PPIH KO2: AAATGGCCCCCGGTAAATAC, ZNF699 (spacer sequence 5’->3’ ZNF699KO1: GCTGGAAAACTTCCAGAACC; ZNF699KO2: CATTCTTACCTAGTGAGGCC) or SETDB1 (spacer sequence 5’->3’ SETDB1 KO1: AAAGCATGTCTTCCCTTCCT; SETDB1 KO2: CAAACCAATGCACCCAGGAA), respectively. For nucleofection, 1 million cells were resuspended in 20 μL of nucleofection buffer P3 (Lonza) and nucleofected with the EA-100 program. At 10 days post-nucleofection, eGFP and tdTomato Cells were mixed 1:1 and incubated with 1 μM indisulam for 4-10 days. Cell populations were assessed by flow-cytometry using a BD LSRFortessa and data were analysed in FlowJo (v10.10.0). Plot was generated by Prism 10; Statistical significance was assessed by two-way repeated-measures ANOVA followed by Šídák’s multiple-comparisons test comparing each knockout with AAVS1 at each time point. *P < 0.05; **P < 0.01; ***P < 0.001; ****P < 0.0001.

### Immunoblotting

For whole cell lysates, cell pellets were washed once with cold PBS, resuspended in 2× Laemmli loading buffer (60 mM Tris-HCl pH 6.8, 10% (v/v) glycerol, 2% (w/v) SDS) at a 1:1 (v/v) ratio with residual PBS, and boiled at 98 °C for 10 min. Protein concentrations were determined using the Qubit Protein Assay Kit (Thermo Fisher Scientific, #Q33212), followed by the addition of β-mercaptoethanol and bromophenol blue from a 20× loading dye stock at a final dilution of 1:20 (v/v). A total of 20 μg of protein per sample was resolved on 4–15% polyacrylamide gels (Bio-Rad) and transferred to PVDF membranes using a dry transfer system (Bio-Rad). Membranes were blocked with 5% (w/v) skim milk in TBS-T for 30 min at room temperature before incubation overnight at 4°C with primary antibodies diluted 1:1000 (anti-ZMAT2, ABclonal; anti-RBM39, Atlas Antibodies (HPA001591); and anti-β-actin, Sigma-Aldrich). The following day, membranes were washed three times for 5 min each with TBS-T and incubated with HRP-conjugated secondary antibodies (Cell Signalling Technology) diluted 1:10000 in 5% (w/v) skim milk for 30 min at room temperature. Membranes were washed 3 times with TBS-T and chemiluminescent signals were developed using ECL substrate prepared by mixing the two components at a 1:1 ratio and imaged using a ChemiDoc Imaging System (Bio-Rad).

### RNA extraction and RNA sequencing

KBM7 cells were nucleofected with sgRNAs targeting AAVS1 or ZMAT2 as described above. Four days after nucleofection, 0.5 × 10^6 cells were seeded per well in 12-well plates containing 1 mL of complete medium. After 16 h, cells were treated with DMSO or 1 μM indisulam for 8 h in technical triplicate before collection. Cell pellets were lysed in TRIzol Reagent (Thermo Fisher Scientific, #15596026), and total RNA was extracted using the Direct-zol RNA Miniprep Kit (Zymo Research, #R2051). RNA concentrations were determined using the Qubit RNA Broad Range Assay Kit (Thermo Fisher Scientific, #Q33265).

For 3′ mRNA sequencing, libraries were prepared using the QuantSeq 3′ mRNA-Seq V2 Library Prep Kit with UMI Second Strand Synthesis Module (Lexogen, #191.96) according to the manufacturer’s instructions. For total RNA sequencing, RNA samples were sent to Novogene for directional library preparation with rRNA depletion and human lncRNA sequencing (WOBI). Libraries from both sequencing approaches were sequenced on an Illumina NovaSeq X Plus platform to generate 150-bp paired-end reads.

### *In-silico* CRISPR screening simulations

All simulations model a pooled screen as a population of 20,000 cells over 21 days of drug selection, evaluated on a grid spaced by the cell-line division time (22 h), partitioned into resistant clones and a non-resistant background pool. Clone fractions Ni(t)/ΣN(t) were rendered as stacked stream plots. Two libraries were simulated under identical initial conditions, differing only in the growth rate assigned to “complete-resistance” clones.

1. Genome-wide library: ten complete-resistance clones with per-clone mean net growth rate drawn from U(1.9, 2.0) day⁻¹ (natural-log basis).
2. Compatible library: the same ten clones assigned a mean growth rate of 0 day⁻¹.

In both scenarios ten “biological-resistance” clones were assigned per-clone mean rates from U(1.6, 1.9) day⁻¹ and the non-resistant pool a mean of 1.4 day⁻¹. Growth was stochastic: at each timepoint the instantaneous rate of each clone was drawn as r ∼ N (clone mean, 0.1), and population size propagated as N_i(t) = N_i(0)·exp(Σ r_i·Δt), a multiplicative random walk. Initial clone sizes N_i(0) were drawn from a discrete uniform distribution on {1,…,5} and the non-resistant pool assigned the remainder of the 20,000 cells. A single realisation per scenario was generated with a fixed seed (123).

The same population model was re-parameterised from the observed screen data rather than from assumed rates. For every gene the MAGeCK positive-selection LFC was converted to a per-day growth advantage over background, a_i = max(LFC_i · ln2 / T_screen, 0), T_screen = 21 days, attributing the observed enrichment entirely to a constant exponential growth differential and flooring depleted genes at zero advantage.

### CRISPR screen analysis

Reads were counted against sgRNA references with mageck count (MAGeCK v0.5.9.5). When comparing the genome-wide Brunello screens against the focused V2 (Brunello–PROTAC V2), both read sets were counted against the same V2 reference, from which sgRNAs targeting the curated ubiquitin–proteasome-system (UPS) genes had already been excluded. Removing these guides at the counting stage rather than physically from the library means the Brunello data are re-analysed post hoc under exactly the V2 gene universe and normalisation base.

Differential sgRNA abundance was tested with mageck test (robust rank aggregation, RRA). Each drug arm was compared with (i) the vehicle (DMSO) arm and (ii) the day-0 (T0) arm. Counts were normalised to non-targeting control sgRNAs (––norm-method control –-control-sgrna), sgRNAs with zero counts in the control sample were discarded (––remove-zero control), and gene-level log2 fold changes were computed as the alpha-trimmed median of the constituent sgRNA LFCs (––gene-lfc-method alphamedian).

#### Gene Ontology

GO Biological Process over-representation was tested with gprofiler2::gost (g:Profiler, Homo sapiens) (v0.2.4). Benjamini–Hochberg FDR correction was applied. Terms with >1,000 annotated genes and a manually curated list of 31 highly generic parent terms were removed. Remaining terms were de-duplicated by hierarchical clustering of the Jaccard string distance between term names (cut height 0.25), retaining the most significant term per cluster.

### RNA-seq analysis

Two datasets were analysed. A 3′-end library set comprising AAVS1 control and two independent ZMAT2 knockouts (ZMAT2KO1, ZMAT2KO2), each treated with indisulam (IND) or vehicle (DMSO), 3 technical replicates per condition, sequenced as 150 bp paired end. A total RNA library set comprising AAVS1 and a single ZMAT2 knockout (ZMAT2KO2), IND or DMSO, three technical replicates per condition, sequenced as 150 bp paired-end.

### 3′-RNA seq preprocessing and counting

For the 3’ RNAseq data, UMIs were extracted from raw reads using umi_tools extract (v1.1.6) and reads were aligned to GRCh38/hg38 using the Rsubread align function (v2.22.1) with default single-end parameters. Gene level read counts were quantified using the featureCounts function (Rsubread v2.22.1) with annot.inbuilt = “hg38” and strandSpecific = 1.

### Total RNA seq Preprocessing and counting

Reads were aligned with STAR v2.7.11b to the human GRCh38 primary assembly genome with GENCODE v45 annotation. A manual two-pass strategy was applied. Briefly, novel splice junctions across all libraries were counted and the alignments were discarded in the first alignment. Novel junctions were pooled and filtered, sites with canonical splice motif and at least 5 supporting reads were kept, whereas 6 unique reads were required for non-canonical splice motifs. In the second pass all libraries were aligned with the filtered novel-junction set. Alignments were then characterized with RSeQC and was inferred to be stranded and reverse as expected. The GENCODE annotation was flattened and overlapping exons were merged with subread (v2.0.6), featureCounts was then performed at the exon level. Transcript abundances were estimated with Salmon (v1.10.2) with 100 bootstrap replicates.

### Gene Level Differential testing

Counts were rounded to integers and analysed with DESeq2 (R v4.5.0, DESeq2 (v1.50.2)), using the design ∼ genotype * treatment, with AAVS1 and DMSO as reference levels. Genes with at least 10 counts in at least 3 samples were retained. Genes were called differentially expressed at Benjamini–Hochberg adjusted-p < 0.05. The equivalent single-KO model (AAVS1 vs ZMAT2KO2) was fitted for the total RNA dataset.

### Gene set enrichment analysis

Pre-ranked GSEA used fgsea v1.36.2 (multilevel; minSize = 10, maxSize = 500, seed 42) against two MSigDB collections obtained with msigdbr (Homo sapiens) v26.1.0: Hallmark (H) and Reactome (C2, CP:REACTOME). Genes were ranked by the DESeq2 Wald statistic; where a symbol mapped to several gene IDs, the ID with the largest absolute statistic was kept.

### Differential testing for exon usage, junction usage, transcript expression and transcript usage

We performed DEU, DTU and DTE with edgeR (v4.8.2). For both quantification levels (transcript-level and exon-level), low count features were removed and TMM normalization was used. The design matrix (∼0 + group) was then fitted and the five contrasts (the interaction, the AAVS1 treatment effect, the KO treatment effect, the KO effect under control, the KO effect under treatment) were tested. We performed differential junction usage testing with ASpli (v2.20.0). The GENCODE v45 annotation was binned and counted using ASpli and tested with all five contrasts.

### Splicing event classification

Alternative splicing was further quantified with rMATS-turbo (v4.3.0) on the STAR-aligned, coordinate-sorted BAMs of the total RNA dataset (12 libraries; AAVS1 and ZMAT2KO2 × DMSO and indisulam, n = 3). rMATS was run against the GENCODE v35 hg38 annotation in paired-end mode (-t paired) with a nominal read length of 150 bp and –-variable-read-length, and a first-strand (dUTP, reverse-stranded) library type (––libType fr-firststrand). rMATS reports five event classes: skipped exon (SE), retained intron (RI), alternative 5′ and 3′ splice sites (A5SS, A3SS) and mutually exclusive exons (MXE).

### Software

Software. Analyses were performed in R v4.5.0 with DESeq2 (v1.50.2), apeglm (v1.32.0), fgsea (v1.36.2), msigdbr (v26.1.0), edgeR (v4.8.2), ASpli (v2.20.0), goseq (v1.62.0), gprofiler2 (v0.2.4), ComplexHeatmap (v2.26.1), circlize (v0.4.18), stringdist (v0.9.17), igraph (v2.2.3), tidygraph (v1.3.1), ggraph (v2.2.2), graphlayouts (v1.2.3), ggstream (v0.1.0), fishplot (v0.5.3), patchwork (v1.3.2), ggpubr (v0.6.3), ggrepel (v0.9.8), Rsubread (v2.22.1) and the tidyverse. Read preprocessing and alignment used umi_tools (v1.1.6), subread (v2.0.6), STAR (v2.7.11b) and Salmon (v1.10.2). MAGeCK v0.5.9.5 was used for CRISPR screen analysis and rMATS-turbo (v4.3.0) for splicing-event classification. Protein association networks used STRING (v12.0) and CORUM (v4.0); flow-cytometry data were analysed in FlowJo (v10.10.0). Figures were generated with ggplot2 and exported as vector PDF.

## Resource Availability

### Lead Contact

Further information and requests for resources and reagents should be directed to and will be fulfilled by the lead contact, Stephin J Vervoort.

### Materials availability

This study did not generate any novel reagents, but desired material or reagents may be requested through the lead contact.

### Data and code availability

Bioinformatics and proteomics datasets generated during this study are available from the NCBI Gene Expression Omnibus (link + GSEA number x).

## Supporting information

FINAL Supplementary Tables 1-18

## Acknowledgments

S.J.V. was supported by a CSL Centenary Fellowship and a Snow Medical Fellowship. This research was supported by the Snow Medical Research Foundation [SMRF2021-SF346]. The funders were not involved in the design of the study, collection, analysis, and interpretation of the data, the writing of this report, or the decision to submit the article for publication. The laboratory of R.F. is supported by The Galbraith Family Charitable Trust, the K & M Foundation for Women, the Betty Deller King Bequest, the Rae Foundation, Denise and Roberto Cappai, John and Tibby Peterson and the Berwick opportunity shop. We greatly thank the Walter and Eliza Hall Institute (WEHI) Genomics and FACS Facilities. We thank Dr Ruifeng Hu for arranging data transfer. We thank the members of the Vervoort lab for critical discussions.

## Author contributions

Conceptualization: S.J.V.; methodology: L.L., O.V. and S.J.V.; software: O.V. and C.Q.W.; validation: L.L.; formal analysis: S.J.V., L.L., O.V., C.Q.W. and M.E.R.; investigation: L.L., O.V. and D.G.; resources: S.J.V.; data curation: O.V., C.Q.W., M.E.R. and R.F.; writing–original draft: S.J.V., O.V. and L.L.; review and editing: L.L., O.V., C.Q.W., D.G., M.E.R., R.F. and S.J.V.; visualization: L.L., O.V. and C.Q.W.; supervision: S.J.V.; project administration: S.J.V.; funding acquisition: S.J.V.

## Declaration of Interests

There are no conflicts of interests to declare.

## Notes

### Competing Interest Statement

The authors have declared no competing interest.

### Summary of Updates

In the previous version PDF file cannot be downloaded, we therfore reuploaded the PDF file,

